# Clock and riboswitch control of *THIC* in tandem are essential for appropriate gauging of TDP levels under light/dark cycles in Arabidopsis

**DOI:** 10.1101/2022.06.20.496815

**Authors:** Zeenat Noordally, Lara Land, Celso Trichtinger, Ivan Dalvit, Mireille de Meyer, Kai Wang, Teresa B. Fitzpatrick

## Abstract

Metabolic homeostasis is regulated by enzyme activities but the importance of regulating their corresponding coenzyme levels is unexplored. The organic coenzyme thiamine diphosphate (TDP) is supplied as needed and controlled by a riboswitch sensing mechanism in plants through the circadian regulated *THIC* gene. Riboswitch disruption leads to loss of time-of-day regulation of *THIC* expression, negatively impacting plant fitness. Pathway precursor balancing combined with enhancing the biosynthesis pathway demonstrate the importance of the riboswitch in gauging TDP levels and indicate that TDP impacts the clock in Arabidopsis. Altering the phase of *THIC* expression to be synchronous with TDP transporters disrupts the precision of the riboswitch suggesting that temporal separation of these processes is important. All defects are bypassed by growing plants under continuous light conditions highlighting the need to control levels of this coenzyme under diel cycles. Thus, consideration of coenzyme homeostasis within the well-studied domain of metabolic homeostasis is highlighted.

## INTRODUCTION

Central metabolism is under strict regulation to maintain homeostasis, requisite for health and fitness of all organisms. Key regulatory enzymes within the core metabolic pathways are assumed to generally deal with maintenance and adjust to address a challenge to homeostasis, the mechanisms of which remain to be fully elucidated (1). A notable challenge exists in plants because the transitions between light and dark periods requires daily reprograming of metabolism, such that the alternation between the energy generating process of photosynthesis during the day and its absence at night can be accommodated (2). Indeed, several metabolites within the central pathways oscillate over diel cycles and are phased to particular times of the day (3), implying diel regulation of enzyme activities. A vast proportion of metabolic enzymes are dependent on organic coenzymes for activity. As the level of loading of the corresponding apoenzyme with the coenzyme dictates at least a pool of catalytic activity, the spatiotemporal provision of coenzyme could be hypothesized to contribute to enzyme driven metabolic homeostasis to be *in sync* with daily light/dark (L/D) cycles (4). However, this has been difficult to track because coenzyme biosynthesis pathways and their regulation need to be unraveled that in turn may provide the requisite tools. Moreover, studies on the significance of keeping coenzyme levels in check are lacking and thus we remain uninformed on a whole facet of metabolic homeostasis and in turn plant health.

A large proportion of organic coenzymes are derived from B vitamins. Recently, the transcripts for biosynthesis and transport of B vitamin derived coenzymes have been shown to be strongly time-of-day regulated in plants (4). One of the most tightly regulated is the coenzyme thiamine diphosphate (TDP) derived from vitamin B_1_ for which biosynthesis and transport are phased to different times of the day (5). Key regulatory enzyme nodes of central metabolism are dependent on TDP as a coenzyme (Figure S1), among which are pyruvate dehydrogenase (glycolysis, Krebs cycle), α-ketoglutarate dehydrogenase (Krebs cycle, amino acid metabolism), transketolase (pentose phosphate pathway and Calvin cycle), and acetolactate synthase (branched chain amino acid synthesis). Interestingly, TDP is short lived (estimated half-life 10 h (6)) and can be destroyed during enzymatic catalysis (7), that drives a high rate of biosynthesis (0.1 nmol h^-1^ g^-1^ in leaves (6)). Biosynthesis involves the suicide enzyme THIAMIN 1 (THI1) (Figure S1) that donates a peptide backbone sulfur atom to the TDP molecule, rendering THI1 non-functional in this context after a single catalytic cycle (8). This results in a remarkably high turnover rate of THI1 (one of the fastest known (9)) that is estimated to consume 2-12% of the maintenance energy budget in plants (10). Therefore, it is thought that luxurious supply of TDP does not occur in plants (11), rather it is made as needed. Coordination of supply to meet demand implies strict monitoring of TDP levels that are intertwined with metabolic requirements. In line with this, TDP levels are monitored in the nucleus by a riboswitch – the only known example in plants (12). The riboswitch operates by alternative splicing in the 3’-UTR of the biosynthesis gene *THIAMIN C* (*THIC*) (13,14) as a function of TDP levels (Figure S1, S2A). Engineering of a dysfunctional riboswitch in Arabidopsis results in plants that are chlorotic and stunted in growth, and is interpreted to be due to excessive levels of TDP that occur in these plants (15). Interestingly, riboswitch mutant plants are unable to adapt to a change in photoperiod (3), suggesting that the riboswitch is particularly important under L/D cycles. However, this has not been explored further and could be important in terms of plant fitness. More generally, the characteristics of TDP and its regulation provides an opportunity to explore the importance of daily surveillance of coenzyme levels.

Here we show that Arabidopsis plants can accommodate enhanced levels of TDP, as long as it is managed appropriately. In particular, time-of-day control of *THIC* transcript levels is critical and should be phased to the evening under L/D cycles managed by the circadian clock and the *THIC* riboswitch in tandem. Disrupting communication and response to TDP status leads to a fitness penalty under L/D cycles that can be bypassed by growing plants under persistent light.

## RESULTS & DISCUSSION

### Enhanced tissue levels of TDP *per se* do not account for the growth defect in Arabidopsis riboswitch mutants

In previous studies, riboswitch mutant lines were generated in the SALK_011114 background (named here as *thiC1-2*) that has a T-DNA insertion in the promoter of *THIC* leading to reduced *THIC* expression. Specifically, *thiC1-2* transgenic lines expressing *THIC* under the control of its own promoter and either the native 3’-UTR (termed here as a *<u>R</u>*esponsive *<u>R</u>*iboswitch (*RR1*)), or a non-functional version of the 3’-UTR carrying an A515G mutation relative to the *THIC* stop codon (termed here as a *<u>N</u>*on-*<u>R</u>*esponsive *<u>R</u>*iboswitch (*NRR1*)) were produced (Figure S2A) (15). Here, we generated a new set of riboswitch mutant lines using the same strategy as before but in the original null allele of *thiC* (SAIL_793_H10) referred to here as *thiC1-1* (Figure S2A), which has an insertion in the third intron of *THIC* (16), to assess its performance. We refer to the new lines as *NRR2* and *RR2*. For comparison with regards to TDP content, we also included engineered lines expressing *THIC* and *THI1* under the control of the *UBQ1* and *CaMV 35S* constitutive promoters and the *UBQ1* and *OCS* terminators, respectively, in wild type Col-0 (referred to here as *TTOE*) (Figure S2A). Similar to *NRR1, TTOE* are reported to have TDP levels above wild type (17). We compared these lines under three different photoperiod regimes, either 8 hr, 12 hr or 16 hr of light. The newly generated *NRR2* lines were chlorotic and had a small leaf lamina with only ca. 50% leaf coverage compared to wild type (Figure 1A) and reduced biomass (Figure 1B). This corroborates previous observations with *NRR1* in the *thiC1-2* background (15) and thus they can be considered morphologically equivalent. However, whereas the phenotype of *NRR* lines persists under all three photoperiods, the *TTOE* lines were morphologically similar to wild type, and with equivalent biomass (Figure 1A, B). To test if a pronounced difference in TDP levels could explain the difference between *NRR1/2* and *TTOE*, we measured TDP content by an established HPLC protocol (5,18). The *NRR1/2* lines showed a significant increase in TDP over wild type (Figure 1C) as reported previously (15), however, the level of TDP in the *TTOE* lines was even higher than the other lines (Figure 1C). Although the levels of thiamine were also significantly increased in *TTOE* lines (Figure S3), these observations suggested that the level of TDP *per se* does not account for the chlorotic stunted growth phenotype observed in *NRR* lines.

**Figure 1.**
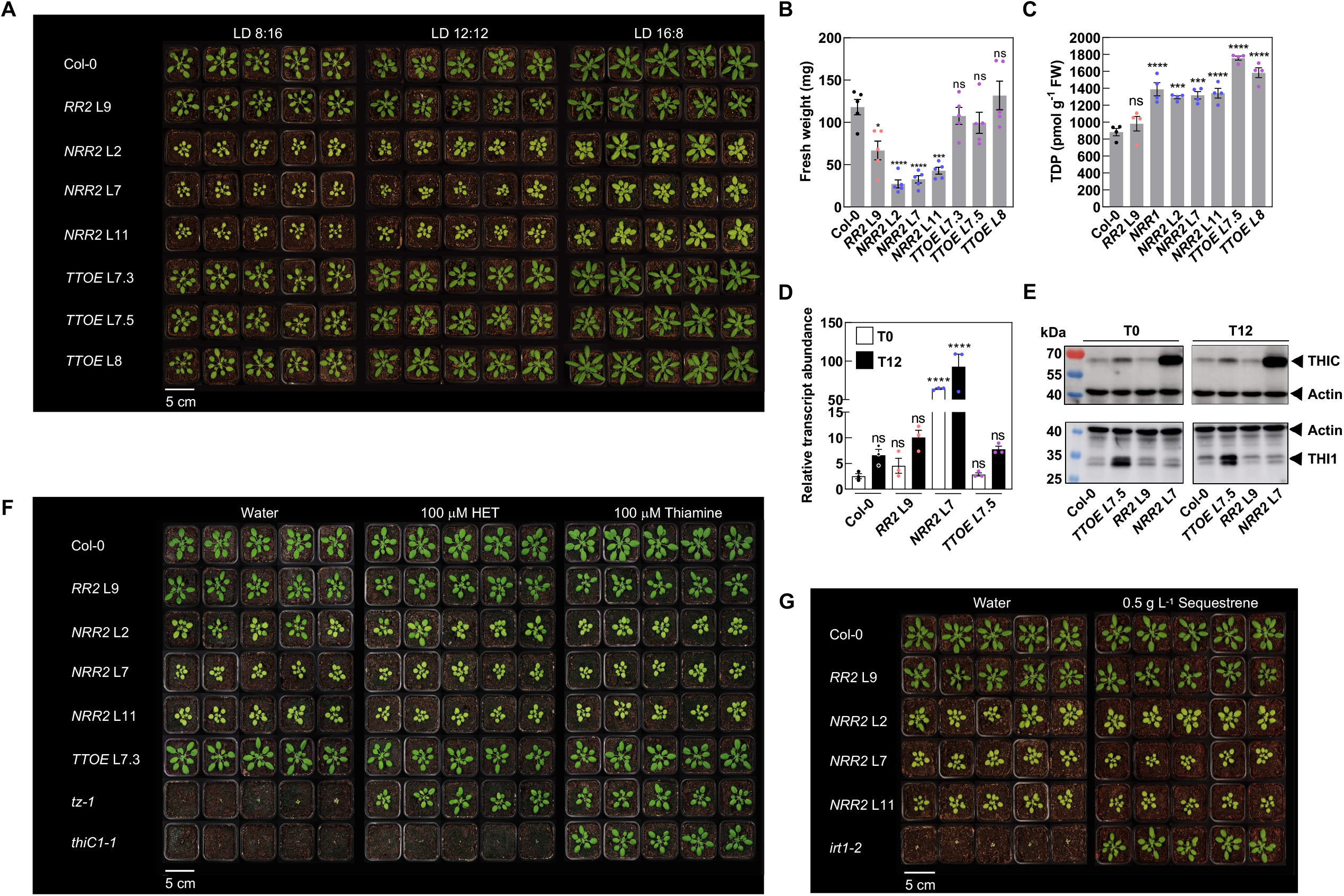
Enhanced TDP content *per se* does not explain the severe phenotype of *THIC* riboswitch mutant plants. (A) Rosette morphology of Col-0 (wild type) Arabidopsis plants compared to transgenic lines in the *thiC1-1* background carrying the wild type (*RR2*) or mutated riboswitch (*NRR2*), and *THIC THI1* overexpressor (*TTOE*) lines in the Col-0 background. Plants were grown on soil under 8, 12 or 16 hr photoperiods (120 to 140 μmol m^-2^ s^-1^ white light) and the corresponding dark period to reach a 24 hr cycle (LD 8:16, LD 12:12, LD 16:8, respectively) and a constant temperature of 20°C. Photos were taken 35 days after germination (DAG) for LD 8:16 and 28 DAG for LD 12:12 and LD 16:8. (B) The fresh weight biomass of shoot material of lines as in (A) grown under LD 12:12 and harvested 28 DAG at 8 hr into the photoperiod. Data represent means of four biological replicates each consisting of a pool of five plants, the standard error of the means (SEM) and one-way ANOVA significance with respect to Col-0. (C) TDP levels of lines as in (B) as well as *NRR1* in the *thiC1-2* background. (D) Expression of *THIC* CDS by RT-qPCR in lines as indicated, grown as in (B) but harvested before the onset of light (T0, white bars) and before the onset of dark (T12, black bars). Transcript levels were normalized to *UBC21*. Data represent means of three biological replicates each consisting of a pool of five plants, the standard error of the means (SEM) and one-way ANOVA significance with respect to either T0 or T12 in Col-0. (E) Immunochemical analyses of THIC and THI1 protein abundance at T0 and T12, with actin as a loading control of plant samples as in (D). Total extracted protein was separated on 12% 37.5:1 polyacrylamide gels. (F) Plants grown as in (B), watered and sprayed twice a week with either water or 100 µM hydroxyethylthiazole (HET) or 100 µM thiamine. The mutant lines *tz-1* (cannot make HET) and *thiC1-1* were used as controls for HET and thiamine rescue, respectively. (G) Plants were grown as in (B), watered and sprayed twice a week with either water or 0.5 g L^-1^ Sequestrene. The *irt1-2* mutant (impaired in iron transport) was used as a control for iron availability. In all cases, significance values are noted as **** for *p* ≤ 0.0001, *** for *p* ≤ 0.001, * for *p* ≤ 0.05 and ns for not significant.

### Balancing thiamine pathway deficits does not salvage improper *THIC* expression levels

In addition to being under riboswitch control, *THIC* transcript levels are strongly influenced by the circadian clock. In particular, an evening element is found in the promoter sequence of *THIC* (Figure S1, S2A) that can bind the morning-expressed Myb-like transcription factors CIRCADIAN CLOCK ASSOCIATED 1 (CCA1) or LATE ELONGATED HYPOCOTYL (LHY) and is reported to repress transcription early in the day (15,19,20). In line with this, transcript levels of *THIC* peak in the evening (5,15). We measured *THIC* transcript abundance in representatives of our set of lines grown for 4 weeks under equinoctial conditions (LD 12:12 cycles), at the onset of light (T 0) and at the onset of dark (T 12). Transcript levels of *THIC* were more abundant at T 12 in wild type (Figure 1D), corroborating earlier reports (5,15). A similar pattern was observed in the *RR* line - the complemented *thiC* carrying an intact riboswitch (Figure 1D). On the other hand, the *NRR* line – *thiC* carrying a dysfunctional riboswitch - showed increased abundance of *THIC* transcript both in the morning and the evening, i.e. constitutive overexpression (Figure 1D). This is despite the transgene carrying the promoter of *THIC* that harbors the evening element (Figure S2A). Thus, clock control of *THIC* expression appears to be lost in the absence of a functional riboswitch. Surprisingly, on the other hand, in *TTOE*, although expression of the *THIC* transgene is under control of the constitutive *UBQ1* promoter and terminator, total *THIC* transcript levels (i.e. endogenous and transgene) were not significantly different to wild type (Figure 1D). Moreover, an increased accumulation at T 12 compared to T 0 was observed similar to wild type (Figure 1D). Thus, time-of-day control of *THIC* transcript levels appears to be intact in *TTOE* which is in the wild type background and thus has a functional endogenous riboswitch. As the level of THIC protein was not examined in previous studies, we also performed immunodetection of THIC in the same samples using an antibody raised against the Arabidopsis protein (16). Interestingly, the level of THIC protein was considerably more abundant in *NRR* compared to any of the other lines, while the level in *TTOE* was only slightly higher than either wild type or *RR* (Figure 1E). It has previously been demonstrated that a balance is achieved (and necessary) between the key enzymes of TDP biosynthesis, THIC and THI1 (Figure S1), to furnish equal provision of the pyrimidine and thiazole heterocycle precursors, respectively, of the TDP molecule (21-23). Thus, we tested the level of THI1 protein in our lines using a peptide antibody raised against the Arabidopsis protein (17). Although, the level of THI1 was increased in *TTOE* as would be expected, there was no significant difference amongst the other lines (Figure 1E). Given the higher abundance of THIC protein in *NRR* and its strong phenotype compared to wild type or *TTOE*, we next tested if supplementation with the thiazole moiety, hydroxyethylthiazole (HET – the product of the THI1 reaction), would compensate for a possible imbalance in the provision of the pyrimidine moiety by THIC. Although, HET supplementation rescued the strong growth defects observed in the *THI1* mutant, *tz-1* (24), it did not improve growth of *NRR* (Figure 1F). Supplementation with thiamine itself rescues both *thi1 (tz-1)* and *thiC* mutant lines as expected, but had no effect on the *NRR* phenotypes (Figure 1F). This suggests that an imbalance in the provision of the thiazole and pyrimidine moieties is not responsible for the phenotype in *NRR*. Of notable interest, is that immunodetection of THI1 results in two bands that may represent two different forms of the protein targeted to either plastids or mitochondria (25). The higher mobility band is relatively less abundant in the evening in all lines and the lower mobility band more abundant in the evening relative to the morning (Figure 1E). This suggests diel regulation of the THI1 protein in contrast to THIC. Next, as the holoenzyme form of THIC carries an iron-sulfur cluster necessary for both stability and activity (16), it is also plausible that overaccumulation of THIC may divest other proteins of this necessary cofactor. Notable amongst iron-sulfur cluster proteins are those of photosynthesis and proteins involved in chlorophyll biosynthesis (26). Moreover chlorosis, as seen in *NRR*, is a diagnostic feature of iron deficiency (27). We therefore tested if iron supplementation could improve the growth of *NRR* on the assumption that assembly of iron-sulfur clusters is not impaired in *NRR* and sulfur is not limiting. Whereas watering with Sequestrene (Fe-EDDHA) restored growth to the *IRON-REGULATED TRANSPORTER1* (*irt1-2*) mutant (28), which in the absence of iron supplementation is stunted in growth and chlorotic, it did not improve growth of *NRR* (Figure 1G).

Taken together, our data show that while TDP levels are increased in both *NRR* and *TTOE*, control of *THIC* expression is different. Time-of-day regulation of *THIC* transcripts is abolished in *NRR* (despite the presence of clock-controlled elements in the promoter of the transgene), while *TTOE* mimics wild type suggesting a complex interaction of the clock and riboswitch in *THIC* transcript control with the riboswitch having a dominant effect.

### Loss of time-of-day *THIC* expression in riboswitch mutants is not due to impaired clock function although TDP impacts clock features

The loss of time-of-day regulation of *THIC* expression drew our attention to the circadian clock and we were next prompted to test if the circadian clock is deregulated in these transgenic lines. We assessed for clock activity by measuring transcript abundance of the reporter genes *CHLOROPHYLL A/B BINDING 2* (*CAB2*) (29,30) by quantitative RT-PCR (qRT-PCR) from plants maintained under equinoctial conditions sampling every 4 hours for 72 hours. An initial visualization of the transcript distributions did not reveal any striking differences between *NRR* or *TTOE* compared to wild type (Figure 2A). Notably, the robust daily rhythm of *CAB2* expression was abolished in the arrhythmic circadian clock mutant of PSEUDO RESPONSE REGULATOR proteins 5, 7 and 9 (*prr5 prr7 prr9*) (31) (Figure S4). We also measured key morning and evening phased components of the genetically encoded circadian clock, *CCA1* and *TIMING OF CAB EXPRESSION1* (*TOC1*), respectively, as well as the day phased constituents *PRR7* and *PRR9* (Figure 2A). In all cases, we assessed period, phase and amplitude using the algorithms from Biodare2 (https://biodare2.ed.ac.uk/) (32) and found that neither period nor phase deviated considerably from wild type, although there were subtle differences in period and phase of *CCA1* and *PRR9* transcripts, respectively, in *NRR* (Figure 2B). The amplitude of *PRR9* was decreased in *NRR* and *TTOE* and that of *PRR7* was decreased in *NRR* (Figure 2B). This implies that the molecular clock is largely operational and near normal in all lines albeit with a reduced amplitude of transcript levels of some components in both *NRR* and *TTOE*. Given that both of the latter have enhanced levels of TDP in common, this suggests that elevated TDP may influence the level of activity of components of the molecular oscillator at least at the transcriptional level. Nonetheless, the data do not support a hypothesis that altered performance of the circadian clock accounts for the stark differences in morphology and diel control of *THIC* expression between *NRR* and *TTOE* and was thus not explored further in the context of this study.

**Figure 2.**
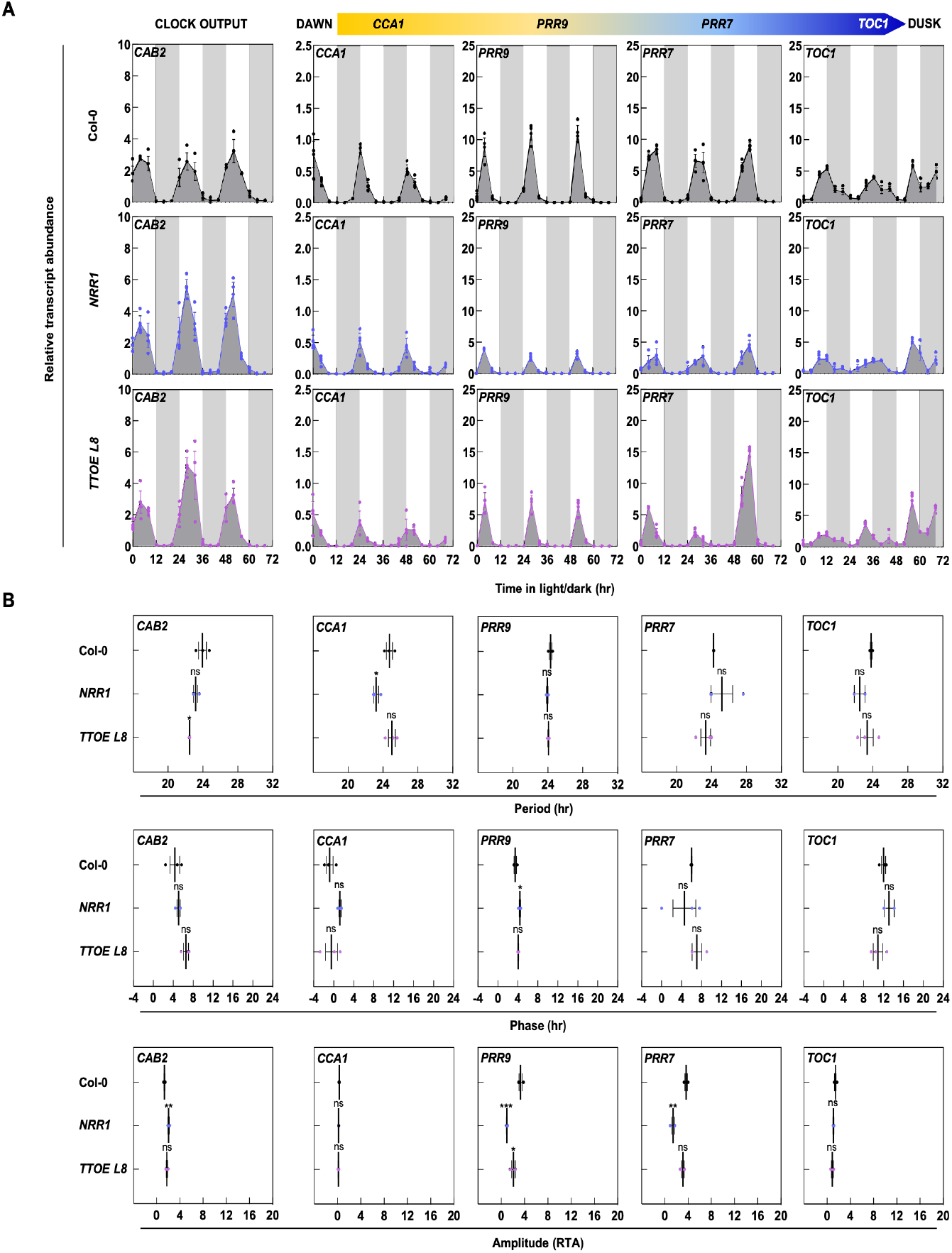
Performance of the clock in lines altered in TDP contents and its sensing. (A) Transcript abundance of clock genes *CCA1, PRR7, PRR9* and *TOC1* that progressively peak from dawn to dusk, respectively, and the clock reporter gene *CAB* by RT-qPCR in Col-0 (wild type, black), mutated riboswitch (*NRR1*, blue) and *THIC THI1* overexpressor (*TTOE*, purple) lines. Plants were grown in culture on 1/2 MS agar plates under a 12 hr photoperiod (120 μmol photons m^−2^ s^−1^ white light) and 12 hr of darkness at a constant temperature of 20 °C. Shoot material was harvested 14 days after germination from pooled seedlings (*n* = 10) every 4 hr at the times indicated. White and gray background bars represent day and night, respectively. Data of three individual experimental replicates of pooled material are shown with error bars representing SE. Transcript levels are relative to *UBC21*. (B) Period, phase, and amplitude (relative transcript abundance, RTA) estimates of gene expression of the Arabidopsis lines in (A) based on the Maximum Entropy Spectral Analysis (MESA) algorithm in BioDare2. Data of three individual experimental replicates are shown with the mean (black line) and error bars representing SEM. Significance values are noted as *** for *p* ≤ 0.001, * for *p* ≤ 0.05 and ns for not significant, calculated by one-way ANOVA followed by Tukey’s multiple comparisons test.

### Sensing of TDP levels is crucial under L/D cycles but not continuous light

As mentioned above, the supply of TDP is presumed to be coincident with time-of-day demand by enzymes dependent on it as coenzyme such that over- or under-supply does not negatively impact plant fitness (6,11). Such tight regulation is postulated to contribute to homeostasis and the natural metabolic reprograming that occurs in plants between day and night cycles (4,15). Support for this comes from the recent observation of daily oscillations of free TDP levels that reflects metabolic differences at the cellular level under L/D cycles and represents the divergence from supply of TDP and the cellular demand by apoenzymes on a daily basis, even though tissue levels of TDP do not change (5). Thus, the riboswitch is expected to be crucial for monitoring cellular levels of TDP and ensuring an appropriate response in line with changes perceived. Therefore, a gauged response from the endogenous riboswitch in *TTOE* might be sufficient to account for the time-of-day regulation of *THIC* expression therein and thus the ability to deal with the enhanced tissue levels of TDP. *NRR* on the other hand would not be able to perceive or transmit alteration of cellular TDP levels due to a non-functional riboswitch. In order to test riboswitch functionality, we monitored splicing of the second intron in the 3’-UTR of *THIC* (referred to as *I*ntron *S*pliced, *IS*) (Figure 3A), which acts as a proxy for the response to TDP levels (5), for 72 hours under equinoctial conditions across all lines. Firstly, in wild type a distinct strong daily rhythm in the abundance of *IS* reflecting the daily oscillation of TDP levels is observed (Figure 3A), corroborating a previous study (5). As expected in *NRR*, overall transcript levels of *IS* are low and no oscillation can be discerned, in line with the impaired splicing (Figure 3A). On the other hand, in *TTOE* there is a strong robust rhythm of *IS* that has an increased amplitude compared to wild type (Figure 3A (note scale difference) and B). As *IS* responds directly to free TDP levels, the increase in *TTOE* suggests perception and response to the higher TDP levels in this line. Splicing of the second intron in the 3’-UTR of *THIC* was confirmed in wild type and *TTOE* as amplification of this region reveals a fragment size of 120 bp (corresponding to splicing of intron 2), while a fragment size of 272 bp is observed for *NRR* corresponding to retention of the second intron (Figure S5A). Furthermore, maintenance of the diel oscillation of *IS* in the arrhythmic *prr5 prr7 prr9* mutant (31) and loss of *IS* oscillation under constant light conditions suggests the *IS* rhythm is responding to TDP as a function of L/D cycles (Figure S5B), corroborating an earlier study (5).

**Figure 3.**
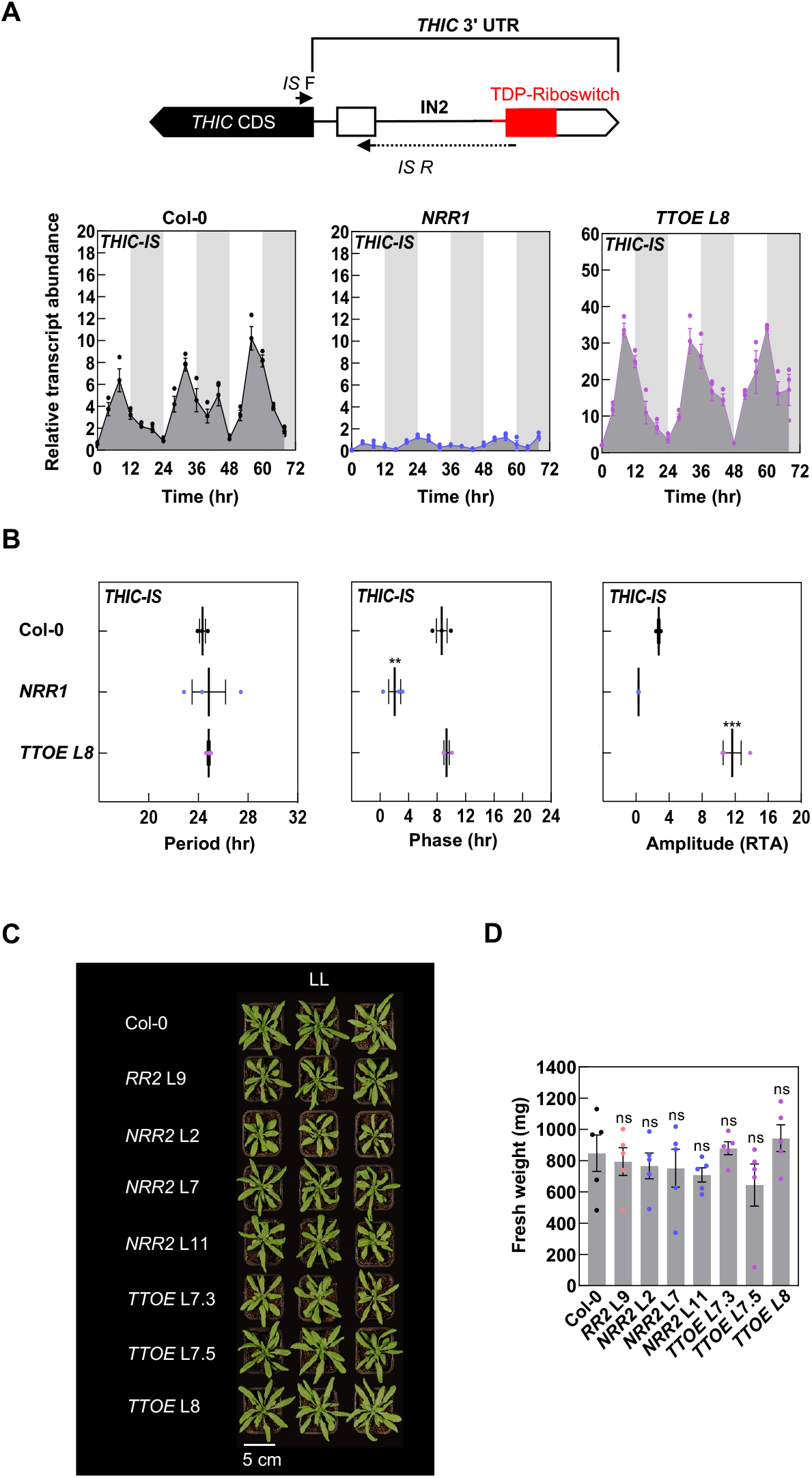
Riboswitch requirement can be bypassed under continuous light. (A) Gene model of part of *THIC*. The coding sequence (CDS; black box), two exons in the 3’-UTR (white boxes), introns as black lines and the riboswitch (red box) are depicted. The abundance of the transcript in which the second intron (IN2) has been spliced (*IS*) is monitored using the *IS* F and *IS* R primers and gauges the response of the riboswitch to TDP. *IS* R (solid black line) spans the spliced site as shown. Transcript abundance of *IS* variants of *THIC* by RT-qPCR. Plants were grown in culture on 1/2 MS agar plates under a 12 hr photoperiod (120 μmol photons m^−2^ s^−1^ white light) and 12 hr of darkness at a constant temperature of 20 °C. Shoot material was harvested 14 days after germination from pooled seedlings (*n* = 10) every 4 hr at the times indicated. White and gray background bars represent day and night, respectively. Data of three individual experimental replicates of pooled material are shown with error bars representing SE. Transcript levels are relative to *UBC21*. (B) Period, phase, and amplitude (relative transcript abundance, RTA) estimates of gene expression of the Arabidopsis lines in (A) based on the Maximum Entropy Spectral Analysis (MESA) algorithm in BioDare2. Data of three individual experimental replicates are shown with the mean (black line) and error bars representing SEM. Significance values are noted as *** for *p* ≤ 0.001, ** for *p* ≤ 0.01, calculated by one-way ANOVA followed by Tukey’s multiple comparisons test. (C) Rosette morphology of Col-0 (wild type) Arabidopsis plants compared to transgenic lines carrying the wild type (*RR*) or mutated riboswitch (*NRR*), or overexpression of *THIC* and *THI1* (*TTOE*). Plants were grown on soil under continuous light (120 to 140 μmol m^-2^ s^-1^ white light) and a constant temperature of 20°C. Photos were taken 28 days after germination. (D) The fresh weight biomass of shoot material of lines as in (C). Data represent means of five biological replicates each consisting of a pool of five plants, error bars represent SE, ns for not significant with respect to Col-0 calculated by one-way ANOVA.

Our data implies that enhanced levels of TDP can be accommodated in Arabidopsis as long as a functional riboswitch is present. With this information in hand, we hypothesized that while gauging of TDP levels by the riboswitch is important for fitness performance under L/D cycles, its requirement can likely be bypassed under continuous light during which a response to adjustment of TDP levels is less important, as daily metabolic reprograming coincident with L/D does not occur. We therefore checked growth of all lines under continuous light and indeed did not observe any morphological difference or significant change in biomass between them (Figure 3C, D), notably for *NRR* which is in stark contrast to its performance under L/D cycles. Thus, the morphological defect resulting from the non-functionality of the riboswitch can be bypassed by growing plants under continuous light.

Taken together, acute sensing of TDP levels through the *THIC* riboswitch suggests its coordination with diel regulation of *THIC* transcript levels. The higher TDP levels appear to be sensed in *TTOE* and are accommodated through increased alternative splicing of (presumably endogenous) *THIC*, which adjusts overall transcript levels to approximate the behavior of wild type. On the other hand, sensing of TDP levels is disrupted in *NRR* and it is ineffective in diel regulation of *THIC* transcript levels. Thus, enhanced levels of TDP can be accommodated in the presence of a functional riboswitch. Nonetheless, how cellular TDP levels are transmitted to the riboswitch remains unknown.

### Correct phasing of *THIC* expression is critical to plant fitness

We postulated that *THIC* expression is coupled to the requirement for TDP under L/D cycles and is gauged by the riboswitch. In parallel, the circadian clock anticipates time-of-day appropriate control, with peak *THIC* expression in the evening. Notably, transcripts for known organellar transport of TDP such as *NUCLEOBASE CATION SYMPORTER1* (*NCS1*) and *THIAMINE PYROPHOSPHATE CARRIER1* (*TPC1*) are temporally separated from those of *THIC* and peak in the morning (5). To test if phasing of *THIC* expression is important and thus temporal separation of TDP biosynthesis and transport, we next designed a strategy to synchronize transcript abundance of both processes to the same time-of-day. As several transporters are implicated in TDP transport (33), the simplest way to engineer this experiment was to phase the peak in expression of the biosynthesis gene *THIC* to the morning (instead of its natural peak in the evening) to be coincident with the peak in abundance of known TDP transporters (around ZT 1-2) (5). To facilitate this, we used two approaches. On the one hand, we placed *THIC* expression under control of the *ACYL CARRIER PROTEIN 4* (*ACP4*) promoter that is phased to ZT 0-2 and shows similar levels of expression to *THIC* in microarray data (Figure S6). For the second approach, we mutated the promoter located evening element (AAATATCT) in *THIC* to the CIRCADIAN CLOCK ASSOCIATED 1-binding site (CBS) element (AAA**A**ATCT) that has previously been shown to change the phase of evening expressed transcripts to the morning (34) (Figure S2B). These constructs *pACP4:THIC* and *pTHIC_EE_CBS:THIC*, both of which carried the 3’-UTR of *THIC* and therefore the riboswitch, were introduced into *thiC1-1* (Figure S2B). Lines homozygous for the respective construct were assessed for time-of-day *THIC* expression. Indeed, the expression levels of *THIC* in *pACP:THIC* almost match those of wild type but are phased to the morning rather than the evening (Figure 4A). Expression levels of *THIC* are phased to the morning rather than the evening also in *pTHIC_EE_CBS:THIC*, although the overall transcript levels are significantly higher than in *pACP:THIC* or wild type, suggesting that this mutation also promotes transcript expression (Figure 4A). Importantly, transcripts of known TDP transporters *NCS1* and *TPC1* were still phased to the morning in these lines and are thus coincident with the expression of *THIC* (Figure S7). Measurement of the tissue TDP content shows that it is not significantly different between the lines (Figure 4B). Next, we measured *IS* in all lines and observed major alterations in the diel rhythms compared to wild type (Figure 4C). Although oscillation can be discerned in *pACP:THIC* the shape of the peaks are much broader and phasing of *IS* is 5.5 hours earlier than wild type (Figure 4C, D). In *pTHIC_EE_CBS:THIC*, the oscillation of *IS* is less robust and the waveform substantially different to wild type (Figure 4C). As TDP levels are similar to wild type in both cases, this suggests that cellular changes under L/D cycles cannot be appropriately gauged and there is less precision in the response of the riboswitch. Such alteration of the *THIC* promoter resulted in plants that were generally chlorotic, especially in the case of *pTHIC_EE_CBS:THIC* and stunted in growth with reduced biomass compared to wild type under L/D cycles (Figure 4E, F). Indeed, these lines are morphologically similar to *NRR*. Remarkably, on the other hand, all lines were indistinguishable under continuous light and in terms of biomass (Figure 4E, F).

**Figure 4.**
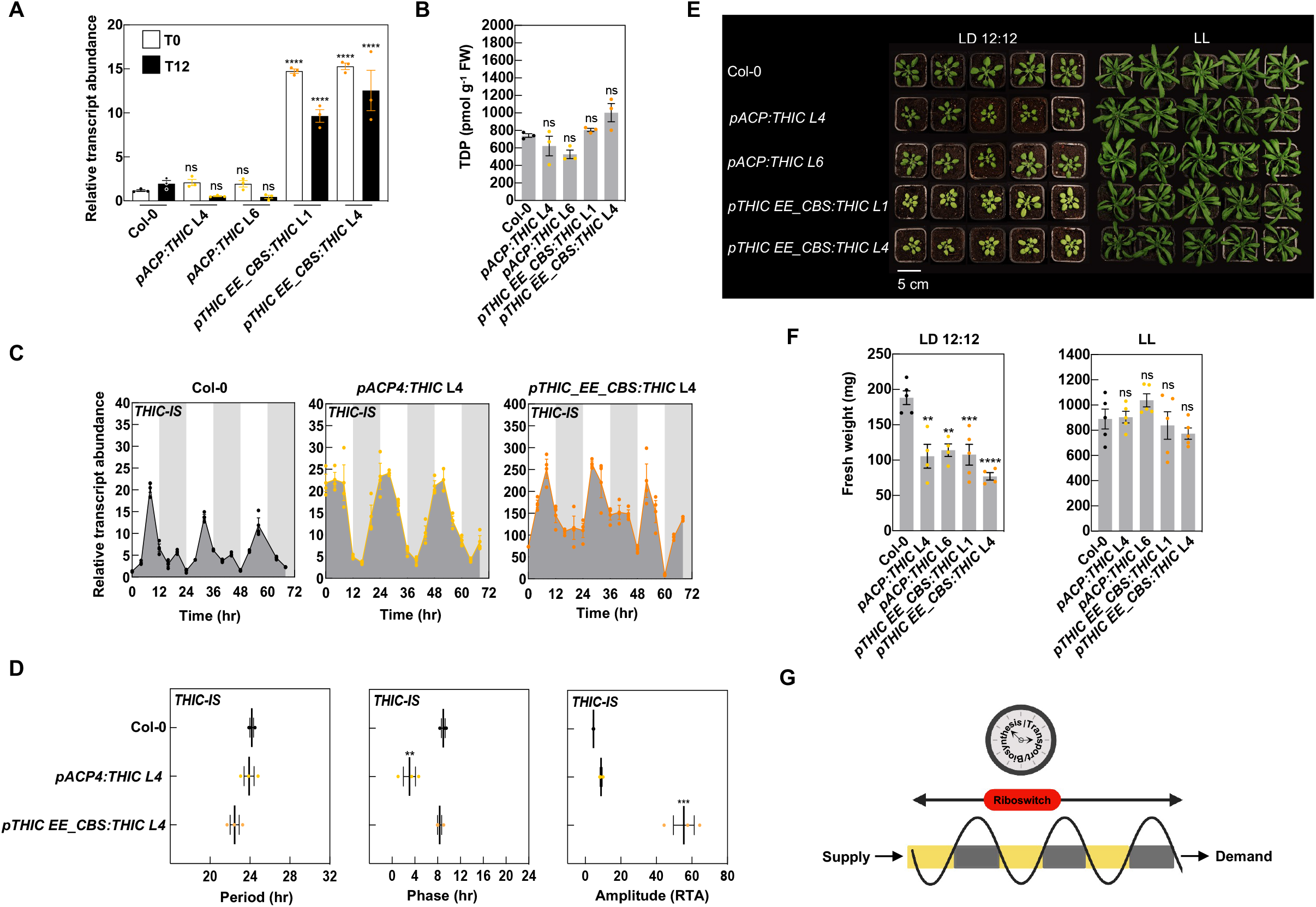
(A) Transcript levels of *THIC* CDS in Col-0 (wild type) compared to transgenic lines in which *THIC* expression is either under control of the promoter of *ACYL CARRIER PROTEIN4* (*ACP4, pACP4:THIC*) or in which the *THIC* promoter based evening element (EE) has been mutated to the *CIRCADIAN CLOCK ASSOCIATED 1-*binding site (CBS, *pTHIC EE_CBS:THIC*). Plants were grown under a 12 hr photoperiod (120 μmol photons m^−2^ s^−1^ white light) and 12 hr of darkness at a constant temperature of 20°C, harvested 28 days after germination (DAG) before the onset of light (T0, white bars) and before the onset of dark (T12, black bars). Transcripts levels were normalized to *UBC21*. Data represent means of three biological replicates each consisting of a pool of five plants, the standard error of the means (SEM) and one-way ANOVA significance with respect to either T0 or T12 in Col-0, where **** = *p* ≤ 0.0001 and ns = not significant are indicated. (B) TDP levels of lines as in (A). (C) Transcript abundance of 3’-UTR second intron spliced (*IS*) variants of *THIC* by RT-qPCR. Plants were grown in culture on 1/2 MS agar plates under a 12 hr photoperiod (120 μmol photons m^−2^ s^−1^ white light) and 12 hr of darkness at a constant temperature of 20°C. Shoot material was harvested 14 DAG from pooled seedlings (*n* = 10) every 4 hr at the times indicated. White and gray background bars represent day and night, respectively. Data of three individual experimental replicates of pooled material are shown with error bars representing SE. Transcript levels are relative to *UBC21*. (D) Period, phase, and amplitude (relative transcript abundance, RTA) estimates of gene expression of the Arabidopsis lines in (C) based on the Maximum Entropy Spectral Analysis (MESA) algorithm in BioDare2. Data of three individual experimental replicates are shown with the mean (black line) and error bars representing SEM. Significance values are noted as *** for *p* ≤ 0.001, ** for *p* ≤ 0.01, calculated by one-way ANOVA followed by Tukey’s multiple comparisons test. (E) Rosette morphology of lines as in (A) grown on soil under a 12 hr photoperiod (120 to 140 μmol m^-2^ s^-1^ white light) and 12 hr of darkness (LD 12:12) or continuous light (LL) and a constant temperature of 20°C. Photos were taken 28 DAG. (F) The fresh weight biomass of shoot material of lines as in (E). Data represent means of four biological replicates each consisting of a pool of five plants, the standard error of the means (SEM) and one-way ANOVA significance with respect to Col-0. (G) Proposed working model for integration of TDP supply and demand over light dark cycles (yellow and dark rectangles, respectively) as a function of the circadian clock control of biosynthesis and transport. In addition, levels of free TDP are gauged by the riboswitch which can act as a rheostat to control levels such that supply and demand are facilitated. In all cases, significance values are noted as **** for *p* ≤ 0.0001, *** for *p* ≤ 0.001, ** for *p* ≤ 0.01 and ns for not significant.

Thus, phasing *THIC* transcript expression to the right time-of-day is important and suggests that temporal separation of TDP biosynthesis and transport by the circadian clock is important for the riboswitch to robustly gauge its response particularly under L/D cycles. As a corollary, coupling between the circadian clock and the riboswitch is critical to appropriately regulate *THIC* transcript levels, which is essential for plant growth and health under environmental cycles.

## CONCLUSIVE COMMENTS

Here a comparison of riboswitch mutants with plants engineered to overexpress the biosynthesis *de novo* pathway genes *THIC* and *THI1* (*TTOE*) allow us to refine the importance of the riboswitch in maintaining TDP levels over L/D cycles. Firstly, both *TTOE* and riboswitch mutants have enhanced levels of TDP. However, in contrast to the chlorotic stunted riboswitch mutants, *TTOE* appear morphologically normal under L/D. Of importance is that riboswitch mutants do not retain diel oscillation of *THIC* expression, rather constitutive enhanced expression is observed, whereas *TTOE* retains diel *THIC* expression. In the riboswitch mutants, the corresponding expected imbalance in thiamine pyrimidine precursor provision cannot be compensated by supplying the thiazole moiety. Similarly, the possible inappropriate hoarding of iron by THIC at the detriment of other proteins is not alleviated by supplying iron. Furthermore, impairment of circadian clock operation does not account for differences between riboswitch mutants and *TTOE*. Rather, *TTOE* appears to adapt to enhanced TDP levels through a gauged response of the riboswitch, as evidenced by increased splicing in the 3’-UTR of *THIC*, that cannot happen in riboswitch mutants. We show that phasing of *THIC* expression to the right time-of-day is important as engineering plants with peak *THIC* transcript phased to a different time of day, in particular to the morning, synchronous with that of TDP transporters, results in unhealthy plants and is linked to imprecision in the response of the riboswitch. We propose that the time-of-day distinct phases of biosynthesis and transporter expression are important for allowing the riboswitch to gauge its response to TDP levels, particularly under L/D cycles (Figure 4G). In support of this hypothesis, riboswitch mutants are indistinguishable from *TTOE* or wild type under continuous light conditions during which metabolic reprograming does not occur and strict monitoring of TDP levels would not be deemed critical. Our data demonstrate that clock and riboswitch control of *THIC* in tandem are essential for appropriate gauging of TDP levels under L/D cycles in Arabidopsis. Given that all transgenic lines perform like wild type under continuous light conditions and the diel rhythm of intron splicing is a function of L/D cycles (5), then communication of TDP status through the riboswitch is highly important under environmental cycles. Under L/D cycles, the maximum level of TDP_free_ is during the day and thus represents the time when supply surpasses demand. Gauging supply during the day may be crucial based on several observations: firstly it was recently suggested that TDP might purposely be curtailed during the day to limit respiratory flux when light serves as a sufficient energy source for plants (11); secondly, sensing of TDP levels is critical for the metabolic reprograming that occurs upon changes in photoperiod and the corresponding change in the timing of the L/D transition (3); and thirdly, in riboswitch disabled plants increased pools of TDP-dependent enzyme activities were concomitant with inappropriate enhanced respiration due to overoxidation of carbon through the TCA cycle leading to starvation before the end of the night (15). While both *NRR* and *TTOE* have enhanced tissue levels of TDP, accurate gauging of levels appears to be in place in the latter as evidenced by the diurnal increased riboswitch activity. This may serve to regulate overall expression of *THIC* in *TTOE* to be similar to wild type. By contrast *NRR* lines are unable to gauge TDP levels and the result is constitutive overexpression of *THIC*. Remarkably, control of *THIC* expression by the circadian clock appears to be suppressed in *NRR*. This suggests an interaction between the riboswitch and the molecular oscillator to be dissected in future studies. Phasing of the riboswitch response relative to the L/D transition is also important as the developmental phenotype of the altered timing of *THIC* expression indicates. This is not merely a consequence of differences in *THIC* expression levels *per se* as the plants behave like wild type under constant environmental conditions. The study highlights the importance of the riboswitch and provides an important rationale for its conservation in plant species and function under L/D cycles. Furthermore, that plants can accommodate enhanced levels of TDP is important for ongoing efforts to improve nutrient content as well as an understanding of its impacts on general plant fitness.

## MATERIALS AND METHODS

### Plant material and growth conditions

*Arabidopsis thaliana* SAIL_793_H10 (N835499) referred to here as *thiC1-1* was from an in house stock (16). SALK_011114 (N511114) referred to here as *thiC1-2* and *tz-1* (N3375, (24)) were obtained from the European Arabidopsis Stock Center. Three independent lines of Arabidopsis carrying the *p35S::THI1 pUBI1::THIC* transgenes, referred to here as *TTOE* (L7.3, L7.5 and L8) were donated by Aymeric Goyer, Oregon State University, U.S.A (17). The *thiC1-2* plants expressing *THIC* under the control of the upstream region relative to the start codon (−1584 to -1 bp) and region downstream of the stop codon including 3’-UTR (2176 to 3581 bp relative to the start codon) with either a Responsive Riboswitch (*RR1*) or Non-Responsive Riboswitch (*NRR1*), which bears a A515G mutation relative to the *THIC* stop codon in the *THIC* 3’-UTR were donated by Asaph Aharoni, Weizmann Institute of Science, Israel (15). Additionally, we generated independent transgenic lines in the *thiC1-1* background expressing similar constructs: the *THIC* coding sequence including the upstream region relative to the start codon (−1584 to -1 bp) and region downstream of the stop codon including 3’-UTR (+2176 to +3581 bp relative to the start codon without or with the A515G mutation relative to the *THIC* stop codon in the *THIC* 3’-UTR) referred to as *RR2* (L2) and *NRR2* (L2, L7 and L11), respectively. For the selection of these lines, T1 seedlings were sprayed with 55 mg/L of glufosinate at 5 days and 8 days after germination (DAG) and selection was maintained at 25 μg ml^-1^ in sterile culture. As the *thiC1-1* T-DNA insertion line and the vector contained the same glufosinate resistance, plant selection was possible due to the thiamine auxotrophy of the *thiC* background. All mutant lines were in the *Arabidopsis thaliana* Col-0 ecotype background (wild type). The altered promoter lines were generated by either substituting the *THIC* upstream region with that of *ACP4* (At4g25050, -688 to +172 bp) or by altering the promoter located evening element (AAATATCT) in *THIC* to the CIRCADIAN CLOCK ASSOCIATED 1-binding site (CBS) element (AAA**A**ATCT). Primers for construct generation and genotyping are given in Table S1. Plants were either grown on soil in CLF Grobank chambers (Plant Climatics, Germany) or in sterile culture in MLR-352 environmental test chambers (Sanyo Electric) under 8 hr L/16 hr D, 12 hr L/12 hr D or 16 hr L/8 hr D photoperiods or continuous light as indicated. Light intensity was 120−140 μmol photons m^−2^ s^−1^, relative humidity was ∼70 %, and temperature was maintained at 20°C. For supplementation experiments, plants on soil were watered and leaves were sprayed either with water (control) or 100 µM 4-methyl-5-thiazole ethanol 98% (HET), or 100 µM thiamine-HCl (Sigma-Aldrich), or watered with 0.5 g/L Sequestrene (Syngenta), as indicated. Seeds grown in sterile culture were surface sterilized, stratified for 4 days at 4°C in the dark, grown on half-strength Murashige and Skoog (MS) medium (35) pH 5.7, without sucrose in 0.55% (w/v) agar (Duchefa) plates. For tissue gene expression, B_1_ vitamer quantification and protein extraction in rosette leaves, shoot material was collected at 28 DAG from plants grown on soil under a 12 hr L/12 hr D photoperiod. Material was harvested in biological triplicates each consisting of a pool of five plants. For time series experiments, plants were grown in sterile culture and split-entrained in diel cycles of 12 hr L/12 hr D or 12 hr D/12 hr L at a light intensity of approximately 120 µmol m^-2^ s^-1^ and at a constant temperature of 20°C. At dawn (0 hr) 13 DAG, plants were either kept in diel cycles of light and dark (diurnal conditions) or transferred to continuous light (circadian free-running conditions). From dawn/subjective dawn at 14 DAG shoot material was harvested at 4 hr intervals, at the times indicated, for three days in biological triplicates, each consisting of material from ten plants. All harvested tissue was immediately frozen in liquid nitrogen and maintained at −80°C until analysis. Frozen tissue was crushed and mixed before ∼50 mg was taken (note that the exact tissue mass for each sample was recorded to normalize the extraction volumes and the data in the HPLC method). The frozen plant material was ground to a fine powder using 2 mm Ø glass beads (Huberlab) and a TissueLyser (Qiagen) for 1 min at maximum frequency. For fresh weight biomass measurements, shoot material was harvested in biological triplicates each consisting of a pool of five plants and weighed on a precision balance (Kern ABS).

### Gene expression analysis by RT-qPCR

Total RNA was extracted using the RNA NucleoSpin Plant kit (Macherey-Nagel) following the manufacturer’s instructions and treated with RNase-free DNase (Macherey-Nagel) to remove trace genomic DNA. Reverse transcription of 1 μg RNA into cDNA was performed using Superscript III reverse transcriptase (Life Technologies) and oligo(dT)15 primers (Promega) according to the manufacturer’s instructions. Fluorescence-based quantitative real time Reverse Transcriptase-PCR analyses were performed in 384-well plates using a QuantStudio5 instrument (Applied Biosystems) and PowerUp SYBR Green master mix (Applied Biosystems). The following amplification program was used: 10 min denaturation at 95°C followed by 40 cycles of 95°C for 15 s and 60°C for 1 min. Data were analyzed using the comparative cycle threshold method (2−ΔCT) and normalized to the reference gene *UBC21* (At5g25760). Primers used are listed in Table S1. Each experiment was performed with at least three biological and three technical replicates.

### Protein extraction and immunochemical analyses

Total protein was extracted from 100 mg of homogenized plant tissue with 300 μl of extraction buffer (50 mM sodium phosphate buffer, pH 7.0, containing 5 mM β-mercaptoethanol, 10 mM EDTA, 0.1% Triton X-100 [v/v] and freshly added 1% [v/v] protease inhibitor cocktail for plant cell extracts (Sigma-Aldrich)). Samples were kept on ice and vortexed for 1 min and centrifuged for 10 min at 16000 *g* at 4°C and the supernatant containing the total protein extract was transferred to a fresh tube. Total protein content was determined using the Bradford method (36), following the manufacturer’s recommendations (AppliChem), and using standard curves derived from known concentrations of bovine serum albumin. Linearity was observed from 0 to 1.5 μg and samples from plant extractions were diluted five-fold with extraction buffer to have a final protein concentration in this range. Twenty μg of total protein extract were loaded and separated by SDS-PAGE on 12% polyacrylamide gels. Migrated proteins were transferred to a nitrocellulose membrane using an iBlot system (Invitrogen) and immunodetection was performed using the SNAP i.d. 2.0 system (Millipore) using a 1:1000 dilution of an *A. thaliana* THIC rabbit primary antibody generated in house (16) or an *A. thaliana* THI1 rabbit primary antibody donated by Aymeric Goyer, Oregon State University, USA (17) and a 1:3000 dilution of goat anti-rabbit IgG horseradish peroxidase (HRP) conjugate secondary antibody (Biorad). Chemiluminescence was detected using WesternBright ECL (Advansta) and captured using an Amersham Imager 680 (GE Healthcare). The membranes were washed with Tris buffered saline containing 0.1% Tween-20 and probed again for protein loading control using a 1:10000 dilution of mouse monoclonal anti-actin (plant) primary antibody (Sigma, A0480) and a 1:3000 dilution of goat anti-mouse IgG HRP conjugate (Biorad).

### B_1_ vitamer quantification by HPLC

B_1_ vitamer content was determined essentially as described in (5) with some modifications. Vitamin B_1_ was extracted at a 1:2 ratio of mg tissue to 1% TCA, typically 50 mg of homogenized plant tissue was resuspended in 100 μl 1% TCA. Samples were vortexed for 10 min at room temperature and centrifuged for 15 min at 16000 *g* at room temperature (ca. 22°C) to remove precipitated proteins. The supernatant was adjusted with 1 M Tris-HCl, pH 9.0, containing 50 mM magnesium chloride (10% v/v). Derivatization of thiamine and its phosphorylated forms to their corresponding thiochromes was achieved by alkaline oxidation, whereby 7 μl of freshly made 46 mM potassium ferricyanide (in 23% sodium hydroxide (w/v)) was added to 56 μl of extracts, vortexed and then incubated in complete darkness for 10 min. Twelve μl of 1 M sodium hydroxide and 25 μl of methanol were then added to the extracts, vortexed and then clarified by centrifugation prior to analysis. Quantification of the B_1_ vitamers was performed in the linear range of a standard curve constructed with known amounts of each vitamer (Applichem) using an Agilent 1260 Infinity series HPLC furnished with a binary pump, an autosampler and a fluorometer (Agilent Technologies). A Hypersil BDS Phenyl 120Å 5 μm guard cartridge 2.0 × 10 mm pre-column (Thermoscientific) was used to protect the column from mobile phase contamination. A Cosmosil π NAP 2.0 ID x 150 mm column (Nacalai Tesque Inc., Kyoto, Japan) was used for stationary phase separation of the B_1_ vitamers. Mobile phase A was 50 mM potassium phosphate buffer, pH 7.2 and mobile phase B was 100% methanol HPLC grade. The following conditions were used: flow, 0.5 mL/min; temperature, 35°C; fluorimetry detection excitation 375 nm, emission 450 nm; PMT gain, 18; peak width, > 0.2 min, 4 s response time, 2.31 Hz; injection volumes, 1 µl and 10 µl; equilibration conditions, 89% buffer A and 11% buffer B. The 20 min running conditions used the following mixtures of buffers A and B; 0 min, 89% buffer A and 11% buffer B; 9.5 min, 68% buffer A and 32% buffer B; 13 min, 40% buffer A and 60% buffer B; 13.5 min, 100% buffer B; 15.5 min, 100% buffer B; 16 min, 89% buffer A and 11% buffer B and 20 min, 89% buffer A and 11% buffer B. Resultant chromatograms were subject to peak integration to quantify B_1_ vitamers present in the plant extracts. Statistical relevance was performed using GraphPad Prism v7.

## CIRCADIAN RHYTHM AND STATISTICAL ANALYSES

All period, phase, and amplitude estimates are based on MESA (Maximum Entropy Spectral Analysis) algorithm in BioDare2 (biodare2.ed.ac.uk; (32)). Statistical analyses were performed in GraphPad Prism v7.

## ACKNOWLEDGMENTS

We gratefully acknowledge the Swiss National Science Foundation (grants 31003A-141117/1 and 31003A_162555/1 to T.B.F.), as well as the University of Geneva for supporting this work. We are grateful to Céline Roux for preliminary HPLC investigations during the course of this study. We thank Asaph Aharoni (Weizmann Institute of Science, Israel) for providing the *NRR1* lines and Aymeric Goyer (Oregon State University, U.S.A.) for the *TTOE* lines.

## AUTHOR CONTRIBUTIONS

T.B.F. conceived the study; Z.N., L.L. and C.T. performed most of the work; K.W. contributed HPLC analysis; I.D. and M. de M. assisted with plant work and analyzed data; T.B.F. wrote the article with assistance from Z.N., L.L. and C.T.

## DECLARATION OF INTERESTS

The authors declare no competing interests.

## Notes

### Competing Interest Statement

The authors have declared no competing interest.

