## Supplemental files for "Clock and riboswitch control of *THIC* in tandem are essential for appropriate gauging of TDP levels under light/dark cycles in Arabidopsis"

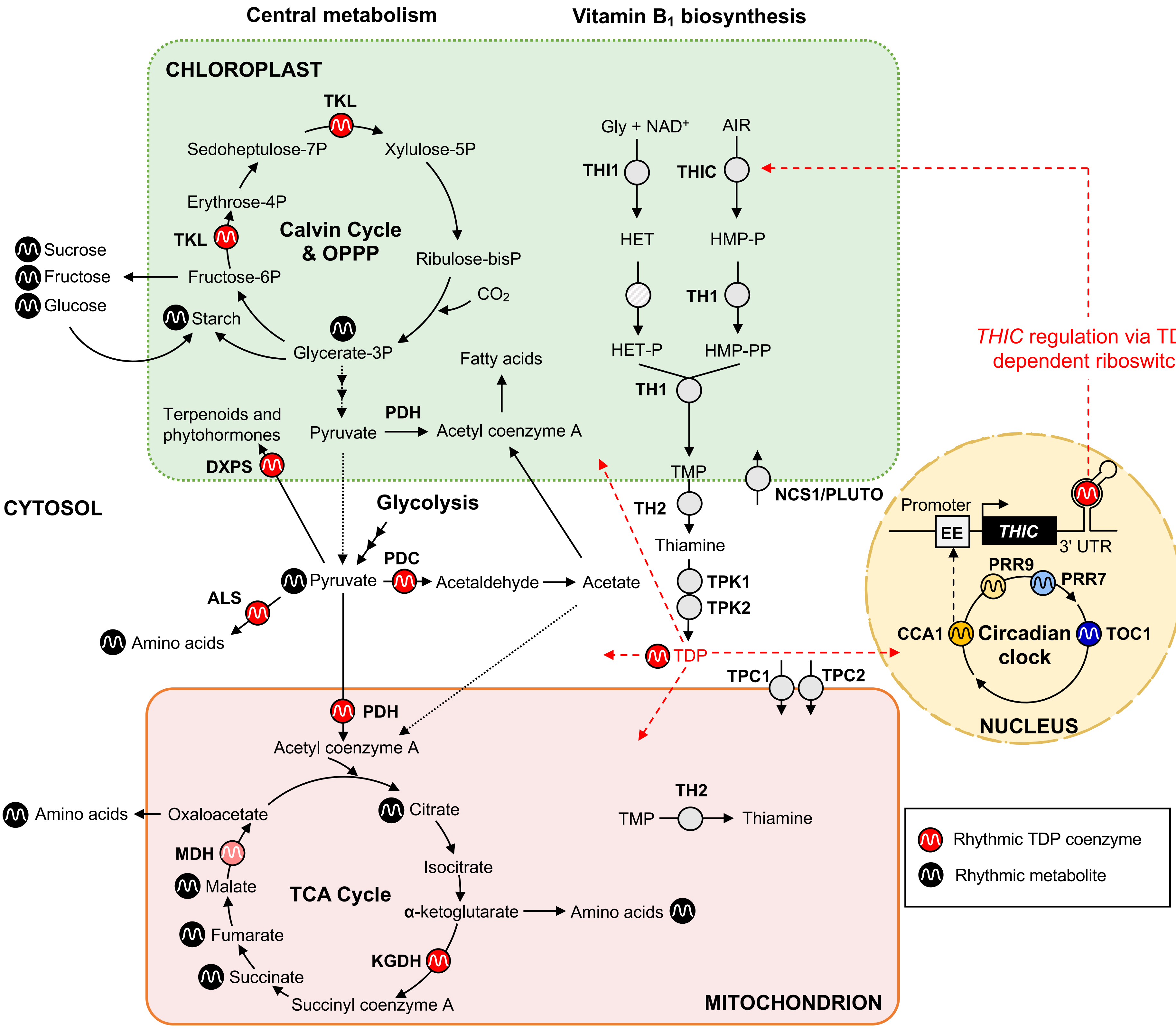


**Figure S1. Plant metabolism is dependent on the vitamin $B_1$ coenzyme TDP.**
An outline of central metabolic pathways is shown, indicating where TDP is required as a coenzyme for enzymes of the Calvin cycle, oxidative pentose phosphate pathway (OPPP), glycolysis and the tricarboxylic acid (TCA) cycle. TDP-dependent metabolic enzymes (red symbols) include transketolase (TKL), pyruvate dehydrogenase (PDH), pyruvate decarboxylase (PDC), 1-deoxy-D-xylulose 5-phosphate synthase (DXPS), acetolactate synthase (ALS), α-ketoglutarate dehydrogenase (KGDH) and potentially malate dehydrogenase (MDH) spread across the chloroplast, cytosol and mitochondria. Where it is known, daily rhythms in abundance of key metabolites are indicated (black symbols) (Gibon et al., 2006; Graf et al., 2010; Bocobza et al., 2013; Sulpice et al., 2014; Rozada-Souza et al., 2019). The vitamin $B_1$ biosynthesis *de novo* pathway is distributed across chloroplasts, cytosol and mitochondria. The thiazole precursor is biosynthesized from glycine and nicotinamide adenine dinucleotide ($NAD^+$) through THI1. The pyrimidine precursor is biosynthesized from aminoimidazole ribotide (AIR) by THIC. The precursors are condensed to form thiamine monophosphate (TMP) by TH1. TMP is dephosphorylated to thiamine by TH2, which is subsequently phosphorylated to the coenzyme form, thiamine diphosphate by TPKs (1&2). TDP relocates from the cytosol to the chloroplasts and mitochondria, where it is required as a coenzyme, and also to the nucleus, where it regulates thiamine biosynthesis *de novo* (red, dashed arrows) through alternative splicing of *THIC* via the riboswitch. Alternative splicing of *THIC* is achieved upon binding of TDP to the riboswitch in the 3'-UTR of *THIC* mRNA (see Figure S2). Additionally, transcription of *THIC*, and much of thiamine biosynthesis and transport, is under the control of the circadian clock. The rhythmic, morning-expressed clock transcription factor CIRCADIAN CLOCK ASSOCIATED 1 (CCA1) binds to the evening element (EE) in the promoter of *THIC*, the release of which allows for transcription of *THIC* at dusk. The transporters through which TDP is relocated serve as gatekeepers for TDP delivery. Other abbreviations used: NUCLEOBASE CATION SYMPORTER 1/PLASTIDIC NUCLEOBASE TRANSPORTER (NCS1/PLUTO), PSEUDO RESPONSE REGULATOR 9 and 7 (PRR9 and PRR7), THIAMINE 1 (THI1), THIAMINE C (THIC), THIAMINE REQUIRING 1 and 2 (TH1 and TH2), THIAMIN PYROPHOSPHOKINASE 1 and 2 (TPK1 and 2), THIAMIN DIPHOSPHATE CARRIER 1 and 2 (TPC1 and TPC2), TIMING OF CAB EXPRESSION 1 (TOC1). Dashed arrows indicate steps that require confirmation.

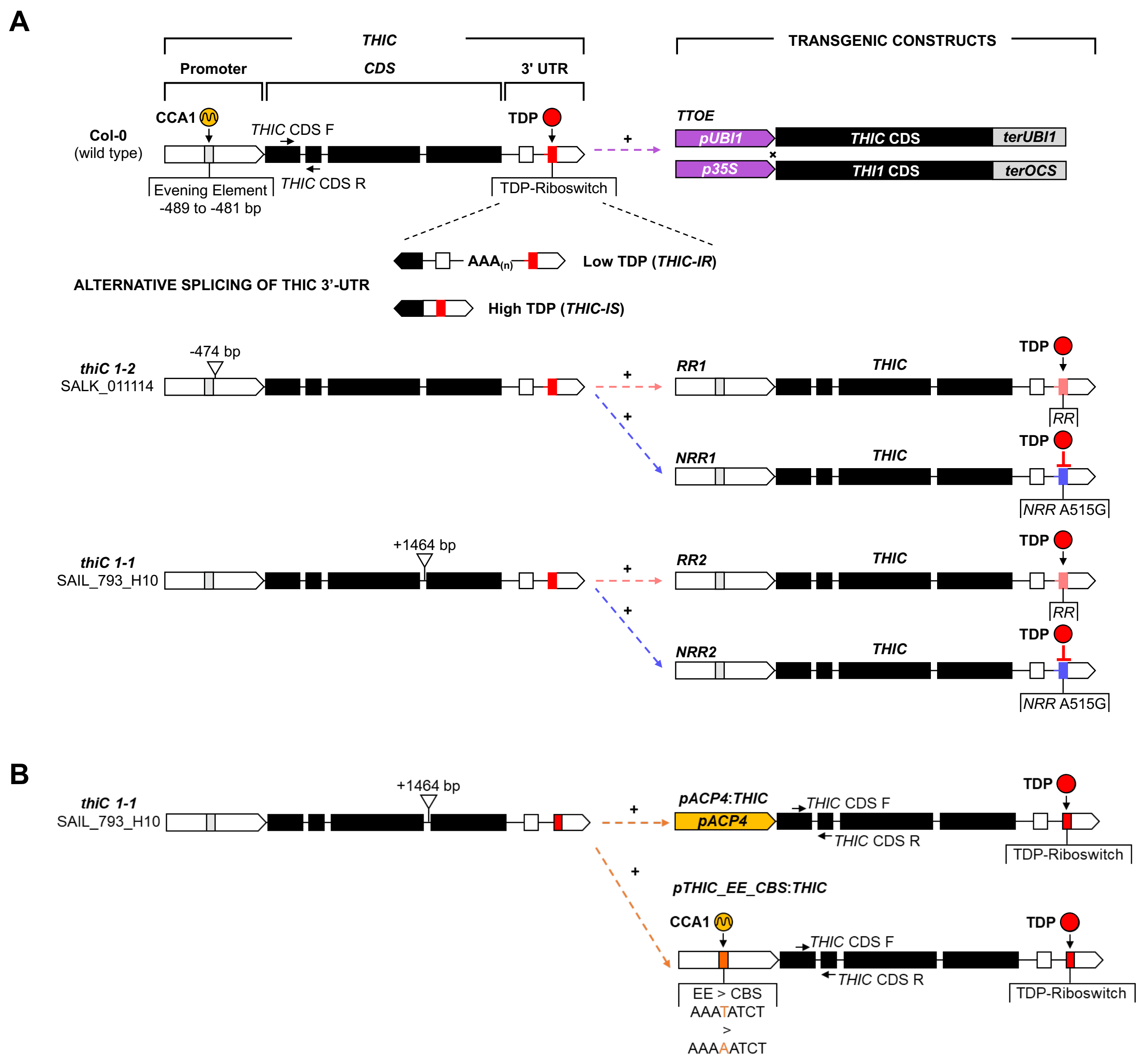


**Figure S2. Illustration of the different transgenic approaches used.**

(A) The gene structure of Col-0 (wild type) *THIC* consists of a promoter region, containing an evening element motif (gray box) recognized by the circadian clock transcription factor CIRCADIAN CLOCK ASSOCIATED 1 (CCA1), and four exons within the coding sequence (CDS; black boxes), as well as two exons in the 3'-UTR (white boxes), introns are depicted as black lines. Arrows indicate forward (F) and reverse (R) primers to amplify *THIC* CDS. The 3'-UTR of *THIC* contains a riboswitch (red box) that can bind free TDP (red circle) that alters splicing of the second intron in the 3'-UTR. When TDP levels are low, this intron that harbors the polyadenylation site ($AAA_{(n)}$) is retained (*THIC IR*). When TDP levels are high the intron is spliced (*THIC IS*). The *THIC THI1 OVEREXPRESSOR (TTOE)* lines express *THIC* and *THI1* in the Col-0 background. The *THIC* construct used harbors the UBIQUITIN1 promoter (*pUB1*, purple box) and terminator (*terUB1*, gray box), while that of *THI1* harbors the promoter of CAULIFLOWER MOSAIC VIRUS 35S (*p35S*) and the OCTOPINE SYNTHASE terminator (*terOCS*), respectively. The section below depicts the position of the T-DNA insertion (inverted triangle) in the *thiC1-2* and *thiC1-1* mutant lines used. These lines were transformed with the constructs shown on the right. Each construct includes the *THIC* upstream region (-1584 to -1 bp from the ATG start codon, where A is +1), *THIC* CDS, and either the Col-0 3'-UTR (+2176 to +3581 bp) annotated as Riboswitch Responsive (*RR*, pink box) or that carrying the A515G mutation that impairs TDP binding and annotated as Non-Riboswitch Responsive (*NRR*, blue box).

(B) *THIC* scheme as in (A). In this case, *thiC1-1* was transformed with the constructs shown on the right that either carry the promoter of *ACYL CARRIER PROTEIN 4*, or a mutation of the evening element (EE) in the upstream region of *THIC* to the *CIRCADIAN CLOCK ASSOCIATED 1-binding site* (CBS). Each of these constructs carry the *THIC* CDS and 3'-UTR that harbors the riboswitch.

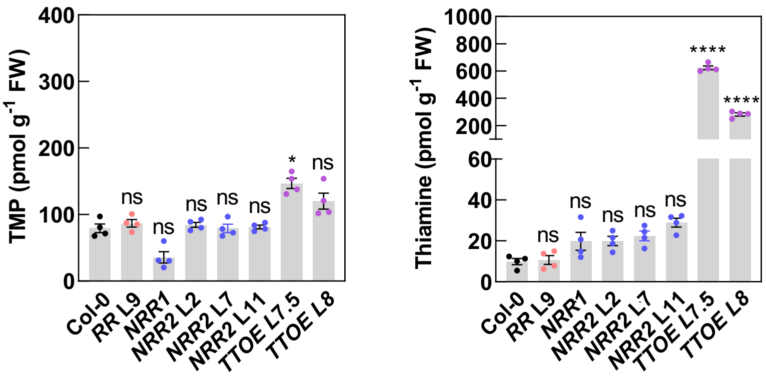


**Figure S3. B vitamer content of transgenic lines compared to Col-0.**
Thiamine monophosphate (TMP) and thiamine content of Col-0 (wild type) Arabidopsis plants compared to transgenic lines in the *thiC1-1* or *thiC1-2* background carrying the wild type (*RR*) or mutated riboswitch (*NRR*), and *THIC THI1 OVEREXPRESSOR* (*TTOE*) lines in the Col-0 background. Plants were grown on soil under a 12 hr photoperiod (120 to 140 µmol $m^{-2}$ $s^{-1}$ white light) and 12 hr of darkness at a constant temperature of 20°C. Shoot material was harvested 28 days after germination at 8 hr into the photoperiod. Data represent means of four biological replicates each consisting of a pool of five plants, the standard error of the means (SEM) and one-way ANOVA significance with respect to Col-0, where **** = $p \leq 0.0001$, * = $p \leq 0.05$ and ns = not significant are indicated.

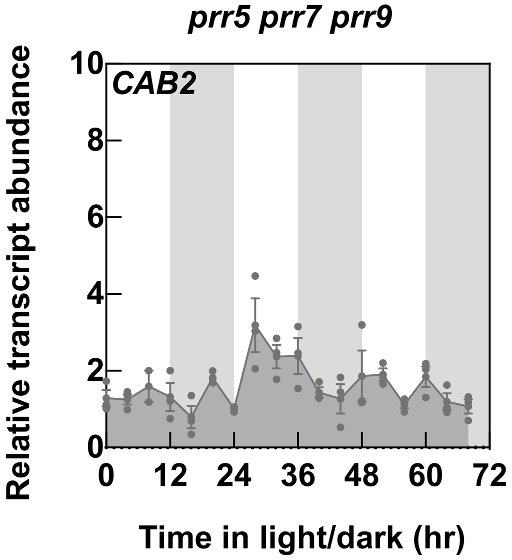


**Figure S4. Rhythmic transcript expression of *CAB* is lost in *prr5 prr7 prr9*.**
Transcript abundance of the clock reporter gene *CHLOROPHYLL A/B BINDING 2* (*CAB2*) by RT-qPCR in the arrhythmic mutant *prr5 prr7 prr9*. Plants were grown in culture on 1/2 MS agar plates under a 12 hr photoperiod (120 µmol photons $m^{-2}$ $s^{-1}$ white light) and 12 hr of darkness at a constant temperature of 20°C. Shoot material was harvested from pooled seedlings ($n = 10$) every 4 hr at the times indicated. White and gray background bars represent day and night, respectively. Data of three individual experimental replicates of pooled material are shown with error bars representing SE. Transcript levels are relative to *UBC21*.

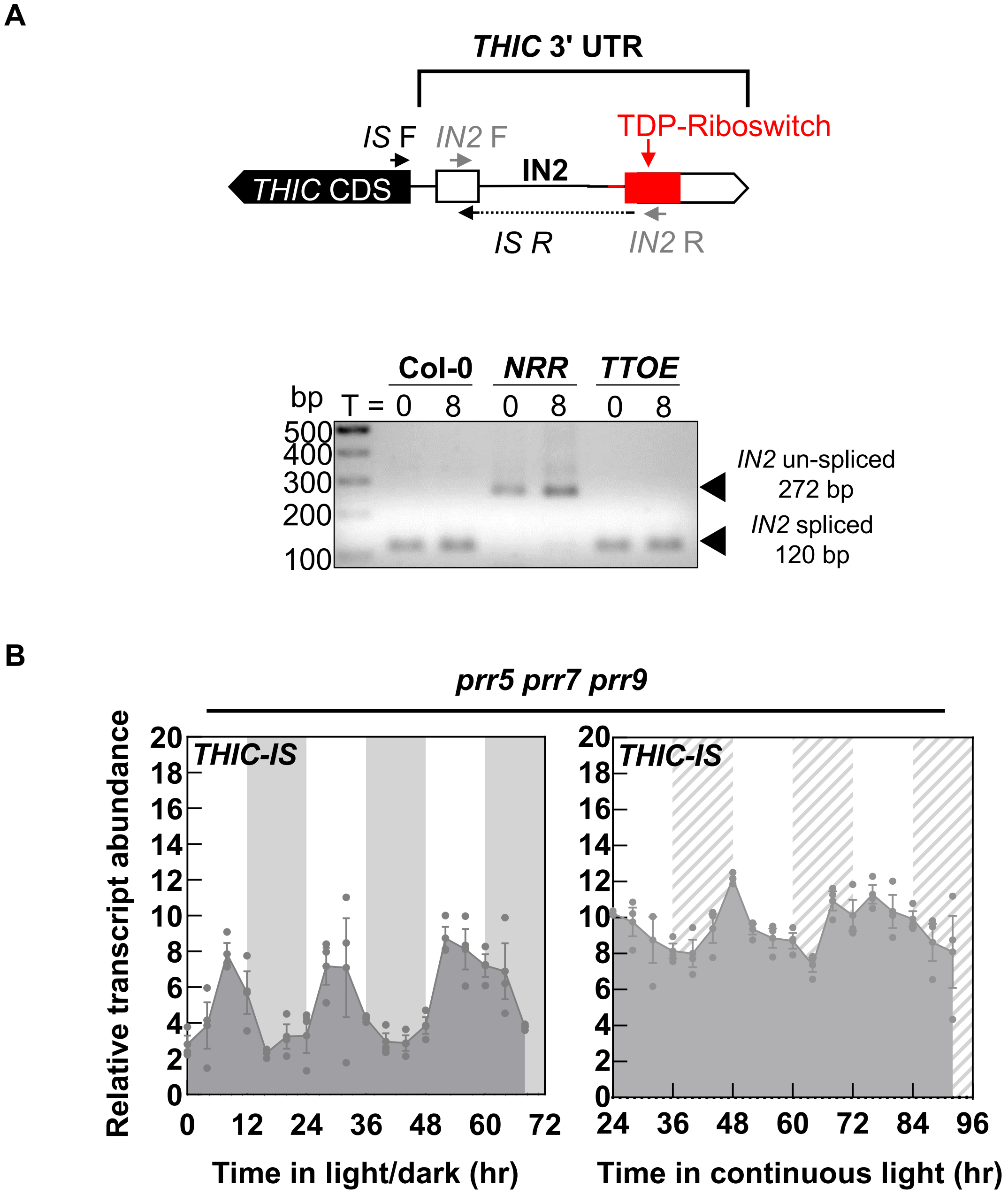


**Figure S5. Assessment of 3'-UTR second intron splicing in *THIC*.**
(A) Gene model of part of *THIC*. The coding sequence (CDS; black box), two exons in the 3'-UTR (white boxes), introns as black lines and the riboswitch (red box) are depicted. The abundance of the transcript in which the second intron (IN2) has been spliced (*IS*) is monitored using the *IS* F and *IS* R primers and gauges the response of the riboswitch to TDP. *IS* R (solid black line) spans the spliced site as shown. The primers *IN2 F* and *IN2 R* are used to amplify the whole region and splicing of IN2 can be deduced from the size. Splicing of intron 2 yields a fragment of 120 bp whereas non-splicing yields a fragment size of 272 bp. The bottom panel shows electrophoresis of PCR amplicons of the 3'-UTR region of *THIC* amplified with *IN2 F* and *IN2 R* (which does not distinguish between variants) on a 2% agarose gel from samples grown for 14 days after germination in equinoctial conditions and harvested at T 0 and T 8 hr.
(B) Transcript abundance of *IS* variants of *THIC* by RT-qPCR in the arrhythmic mutant *prr5 prr7 prr9* under equinoctial light dark or continuous light cycles. White and dark gray areas represent light and dark in equinoctial conditions, whereas white and hatched gray areas represent subjective day and night in continuous light conditions. Plants were grown in culture on ½ MS agar plates and entrained for 13 days in equinoctial conditions (12 hr photoperiod with 120 µmol photons $m^{-2}$ $s^{-1}$ and 12 hr of darkness at 20°C) and either transferred to constant light or retained in equinoctial conditions. Shoot material was harvested over 3 days at 4 hr intervals from seedlings ($n = 10$) at the times indicated. Data of three individual experimental replicates of pooled material are shown with error bars representing SE. Transcript levels are relative to *UBC21*.

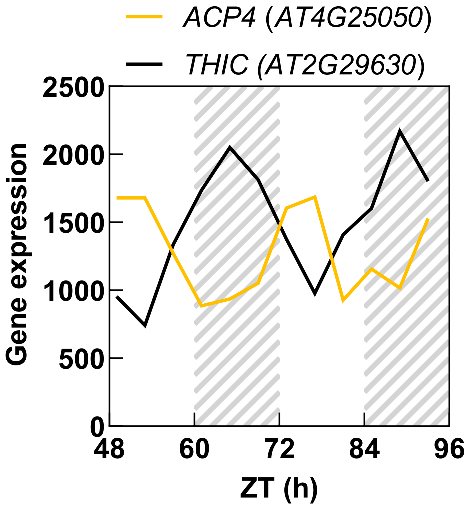


**Figure S6. Anti-phasic expression of *ACP4* relative to *THIC*.**
Transcript levels of *ACYL CARRIER PROTEIN4* (*ACP4*) compared to *THIC* taken from the diurnal database (Mockler, T.C. 2007). The condition 9-day-old Col-0 seedlings grown on ½ MS agar, no sucrose, in continuous light (100 µmol photons $m^{-2}$ $s^{-1}$) at 22°C (LL-LLHC) was used to extract data.

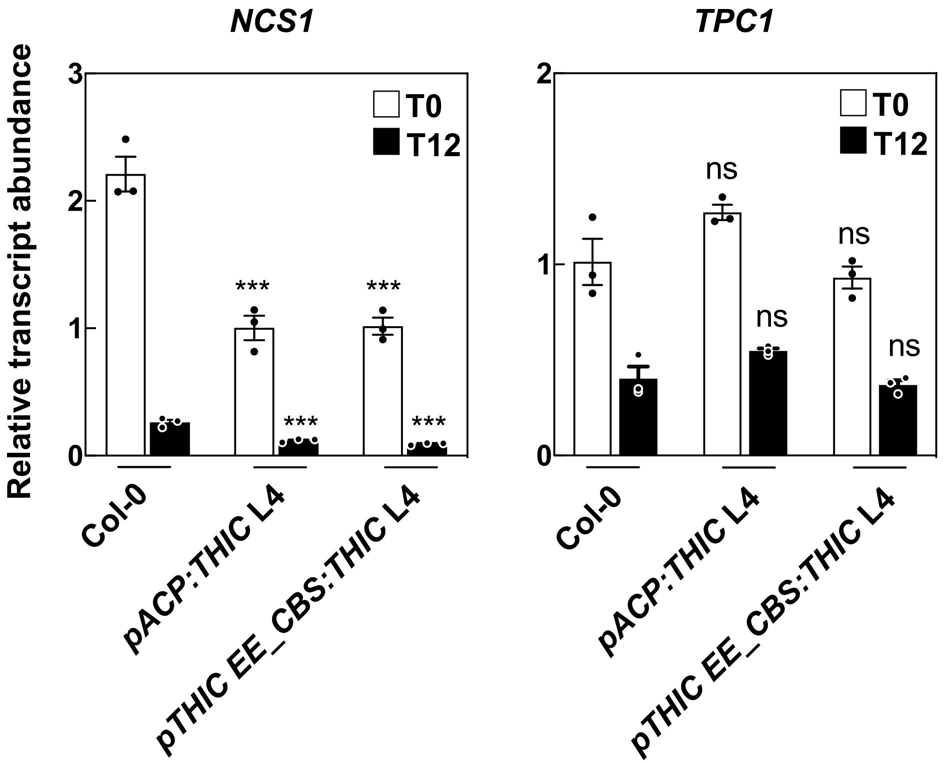


**Figure S7. TDP transporters are phased to the morning.**

Expression of transporters *NCS1* and *TPC1* by RT-qPCR in lines as indicated harvested before the onset of light (T0, white bars) and before the onset of dark (T12, black bars). Transcript levels were normalized to *UBC21*. Plants were grown on soil under a 12 hr photoperiod (120 to 140 µmol $m^{-2}$ $s^{-1}$ white light) and 12 hr dark and a constant temperature of 20°C. Data represent means of three biological replicates each consisting of a pool of five plants, the standard error of the means (SEM) and one-way ANOVA significance with respect to either T0 or T12 in Col-0.

In all cases, significance values are noted as *** for $p \leq 0.001$ and ns for not significant.

**Table S1.** List of oligonucleotides used in this study.

| **Name** | **Locus ID** | **Direction** | **Sequence (5′ – 3′)** | **Purpose** |
|---|---|---|---|---|
| *UBC21* | At5g25760 | Forward | TAGCATTGATGGCTCATCCTGA | RT-qPCR normalization gene |
| *UBC21* | At5g25760 | Reverse | TTGTGCCATTGAATTGAACCC | RT-qPCR normalization gene |
| *THIC CDS* | At2g29630 | Forward | CCATCTTTTGAAGAATGCTTTCCT | RT-qPCR |
| *THIC CDS* | At2g29630 | Reverse | GAACACGACGAAAGGGAACTTT | RT-qPCR |
| *THIC IN2* | At2g29630 | Forward | CTTGGTGCCTGTTGGACTATACC | RT-qPCR |
| *THIC IN2* | At2g29630 | Reverse | TCAGGTTCAAAGGGACTTTCTCA | RT-qPCR |
| *THIC IS* | At2g29630 | Forward | GCTATGTCAAAGCTGCTCAGAA | RT-qPCR |
| *THIC IS* | At2g29630 | Reverse | CAAGCACCCCAACAGTTTGT | RT-qPCR |
| *CCA1* | At2g46830 | Forward | GCACTTTCCGCGAGTTCTTG | RT-qPCR |
| *CCA1* | At2g46830 | Reverse | TGACTCCTTTCTTACCCTGTTATTCTG | RT-qPCR |
| *PRR9* | At2g46790 | Forward | TGCGTCGGCCTTCTCAAGATTTG | RT-qPCR |
| *PRR9* | At2g46790 | Reverse | TGGTGTCTTTGGCTCACCTGAAGT | RT-qPCR |
| *PRR7* | At5g02810 | Forward | CCACGAGCGGTATCTCTATGG | RT-qPCR |
| *PRR7* | At5g02810 | Reverse | ACTGATTACTTGGAAACTCAGGGTTAG | RT-qPCR |
| *TOC1* | At5g61380 | Forward | TCTTCGCAGAATCCCTGTGAT | RT-qPCR |
| *TOC1* | At5g61380 | Reverse | GCTGCACCTAGCTTCAAGCA | RT-qPCR |
| *CAB2* | At1g29920 | Forward | TCAATCTTTTGAATTCGAGTGAGA | RT-qPCR |
| *CAB2* | At1g29920 | Reverse | TCCACCACAAACACAAACCTAC | RT-qPCR |
| *NCS1* | At5g03555 | Forward | CGTGTTTCAGCCATGGAGATTGC | RT-qPCR |
| *NCS1* | At5g03555 | Reverse | AAGCGCTGAGTACCCTATGAGC | RT-qPCR |
| *TPC1* | At3g21390 | Forward | TATGCTGGTCTGCAGTTT | RT-qPCR |
| *TPC1* | At3g21390 | Reverse | GCTTGATGAAGAAGATCGGT | RT-qPCR |
| *pCAMBIA_KpnI_pTHIC* | At2g29630 | Forward | GTATCTCCGTCAGGTACCTCTTCTCCTTCTAG | Cloning *KpnI* restriction site in *pCAMBIA_THIC* vector |
| *pCAMBIA_KpnI_pTHIC* | At2g29630 | Reverse | CTAGAAGGAGAAGAGGTACCTGACGGAGATAC | Cloning *KpnI* restriction site in *pCAMBIA_THIC* vector |
| *pCAMBIA_pTHIC_NcoI* | At2g29630 | Forward | CGTTTGTCTCCATGGAATGGCTGCTTC | Cloning *NcoI* restriction site in *pCAMBIA_THIC* vector |

| | | | | |
|---|---|---|---|---|
| *pCAMBIA_pTHIC_NcoI* | At2g29630 | Reverse | GAAGCAGCCATTCCATGGAGACAAACG | Cloning *NcoI* restriction site in *pCAMBIA_THIC* vector |
| *pCAMBIA_KpnI_pACP4* | At4g25050 | Forward | CGGGGTACCGATTTTCTTACATTTTATATCATACAATTCAATTCC | Cloning *pACP4* promoter into *pCAMBIA_THIC* vector |
| *pCAMBIA_pACP4_XmaI* | At4g25050 | Reverse | TCCCCCCGGGTTGAAGGAGATGAAGCTCAATACAC | Cloning *pACP4* promoter into *pCAMBIA_THIC* vector |
| *pTHIC_EEΔCBS* | At2g29630 | Forward | CCAATTTTCGACAAAAAATCTGAGAAAGAGGAC | Mutagenesis |
| *pTHIC_EEΔCBS* | At2g29630 | Reverse | GTCCTCTTTCTCAGATTTTTTGTCGAAAATTGG | Mutagenesis |
